# Bird interactions with drones, from individuals to large colonies

**DOI:** 10.1101/109926

**Authors:** Mitchell Lyons, Kate Brandis, Corey Callaghan, Justin McCann, Charlotte Mills, Sharon Ryall, Richard Kingsford

**Affiliations:** Centre for Ecosystem Science, School of Biological, Earth and Environmental Sciences, UNSW Australia, Sydney

**Keywords:** drones, UAV, UAS, birds, breeding, waterbirds, raptors, magpies

## Abstract

Drones are rapidly becoming a key part of the toolkit for a range of scientific disciplines, as well as a range of management and commercial applications. This presents a number of challenges in context of how drone use might impact nearby wildlife. Interactions between birds and drones naturally come to mind, since they share the airspace. This paper details initial findings on the interactions between drones and birds for a range of waterbird, passerine and raptor species, across of a range of scientific applications and natural environments. The primary aims of this paper are to provide guidance for those planning or undertaking drone monitoring exercises, as well as provide direction for future research into safe and effective monitoring with drones. Our study sites we all located within Australia and spanned a range of arid, semi-arid, dunefield, floodplain, wetland, woodland, forest, coastal heath and urban environments. We particularly focus on behavioral changes towards drones during breeding season, interactions with raptors, and effects on nesting birds in large colonies – three areas yet to be explored in published literature. In over 70 hours of flight, there were no incidents with birds. Although some aggressive behavior was encountered from solitary breeding birds. Several large breeding bird colonies were surveyed, and included in our observations is monitoring and counting of nests in a colony of over 200,000 Straw-necked Ibis, the largest drone-based bird monitoring exercise to date. In addition to providing observations of interactions with specific bird species, we recommend procedures for flight planning, safe flying and avoidance. This paper also provides a basis for a number of critical and emerging areas of research into bird-drone interactions, most notably, territorial breeding birds, safety around large raptors, and the effect of drones on the behaviour of birds in large breeding colonies.

## Introduction

Unmanned aerial vehicles (hereafter drones), with their varied applications and general affordability, are increasingly used in ecological research and monitoring. Surveying birds from the air has many benefits (Kingsford and Porter, 2009). Use of drones in this context has a surprisingly long history (Abd-Elrahman et al., 2005; Chabot and Francis, 2016). Whilst application to avian research and management is relatively limited compared to other disciplines, it is gaining momentum. Current research spans a range of topics, including ethical guidelines (Vas et al., 2015), recreating environmental data input from bird flight paths (Rodríguez et al., 2012), monitoring nesting status (Weissensteiner et al., 2015), and both manual and automated detection routines for groups of birds and nest counts (Chabot and Bird, 2012; Chabot and Francis, 2016; Sardà-Palomera et al., 2012; Hodgson et al., 2016; Descamps et al., 2011; Trathan, 2004).

There are a range of challenges related to collection of data using drones, and a major component of this is interaction with nearby wildlife (Lambertucci et al., 2015). Naturally, birds are of great interest, given that they share the airspace. Research has only just begun in exploring these interactions (Vas et al., 2015), identifying a considerable knowledge gap in context of the diversity of bird species and how they interact with drones. In the context of drones, there is currently no literature on behavioral changes with breeding status, interactions with raptors, and effects on nesting birds in large colonies. Most parts of the world also have very little information about interactions with drones and local bird species. In this paper we provide some initial findings and guidelines to address some of these knowledge gaps. Drawing observations from over 70 hours of flight, we detail bird-drone interactions across a wide range of environments.

We particularly focus on observations of birds during their breeding season, when nesting birds are more likely to be susceptible to disruption (Lambertucci et al., 2015). During the breeding season, drones can be particularly hazardous for the birds, given potential large congregations and territorial aggression. Of particular interest are our observations while monitoring several large breeding waterbird colonies; one colony contained at least 100,000 nests. To date, the largest reported colony of birds monitored via a drone is a penguin colony of 11,000 (Trathan, 2004). We also report a number of interactions with raptors. Further to detailing the interactions with various bird species, we also provide some recommendations for safe flying and avoidance. This paper also provides the first comprehensive report of bird-drone interactions in Australia. The primary aim of this paper is to provide a basis for further research into bird-drone interactions, and to help readers in planning and safely executing monitoring work with the use of drones.

## Material and methods

### Study locations and monitoring details

Our study locations are within predominantly within eastern Australia but we focus on bird species that have a continental distribution (Fig. 1). The cluster of sites around Sydney were at various National Parks and urban greenspaces. The remaining sites were spread across a range of environments including arid and semi-arid floodplains, shrublands and dunefields, as well as permanent wetlands. Drone use spanned a range of survey planning and environmental monitoring activities. Table provides study site details, including the purpose of drone use and flight characteristics. Exact locations are not provided due to sensitivity for breeding birds, but are available from authors on request. Except for the Ibis colonies, bird observations were made incidental to normal drone operation activities. For the Ibis colonies, we conducted more systematic observations, which are detailed below. The main drone used for monitoring at all sites was a *DJI Phantom 3 Professional* quad-copter. Additionally, a *Sensefly eBee* fixed-wing and a *DJI S900* hexa-copter was also flown at some sites.

**Figure 1:**
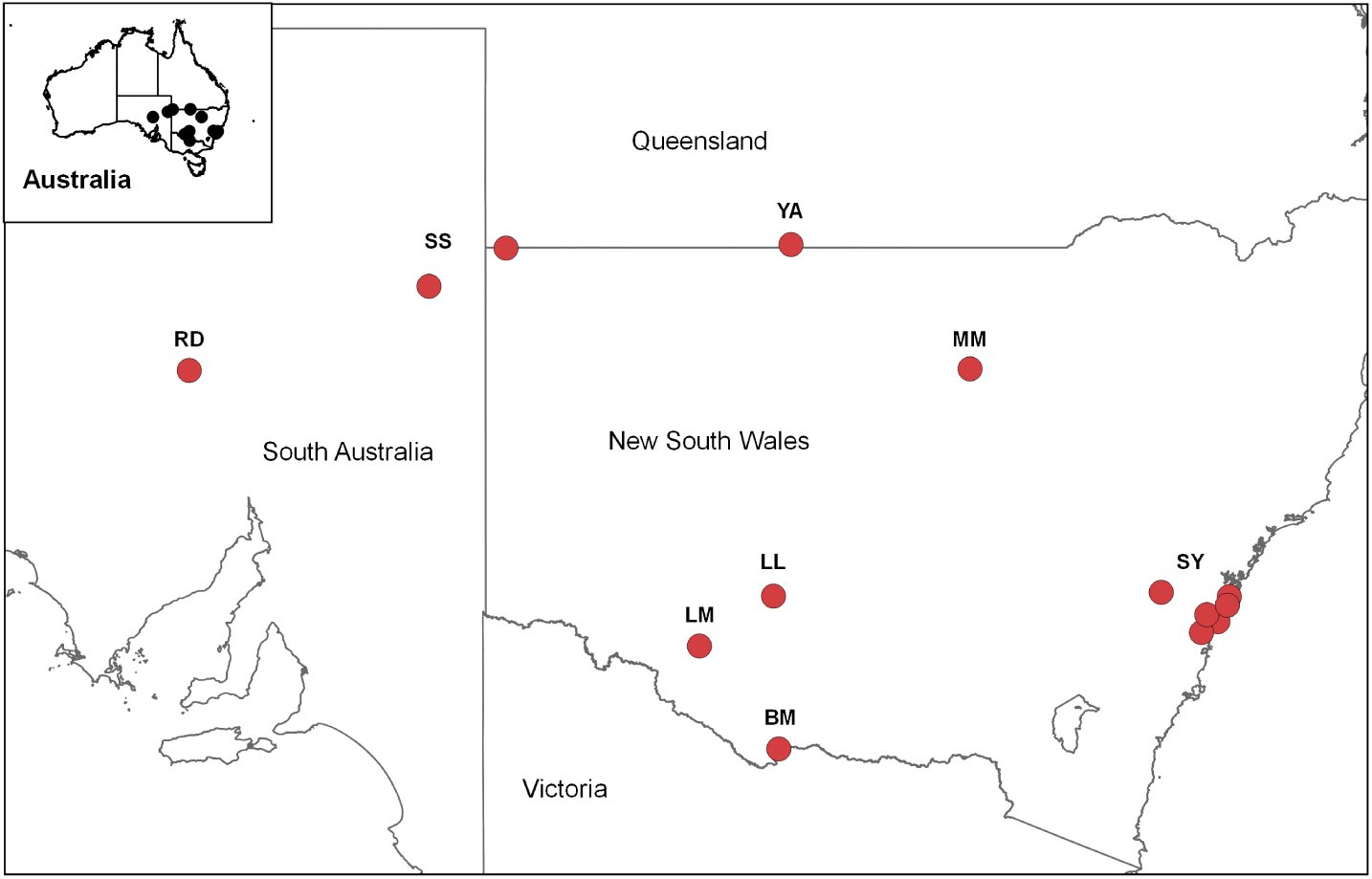
Map showing study locations for this paper. BM=Barmah-Millewa; LL=Lower Lachlan; LM=Lower Murrum-bidgee; MM=Macquarie Marshes; RD=Roxby Downs; SS=Sturt and Strzelecki; SY=Sydney Basin/City. See Table for more details.

### General flight details

The main purpose for drone use at most of the study sites was to acquire imagery to generate orthorectified mosaics and related 3D model products. This typically involved flying parallel flight lines at speeds between 5 to 10 m/s. To acquire sufficient image overlap for processing, flight lines were typically 20 to 100 m apart depending on flying height. For example, flying at 100 m above take off (ATO), flight lines were around 100 m apart, whereas at 20 m ATO, flight lines were around 20 m apart. As an example, one of the Lower Lachlan River surveys covered an approximately circular area of around 7 km^2^ and we flew 34 individual flight transects at 100 m ATO. As most of the monitoring was in wet, muddy or dusty environments, the *DJI Phantom 3 Professional* was predominantly used, as it is relatively affordable. For example, the bird colonies were entirely under water, so failure or emergency landing would result in loss of the drone. Incidentally, all terrain vehicles provide a good platform for take off in a range of environments (Fig. 2).

**Figure 2:**
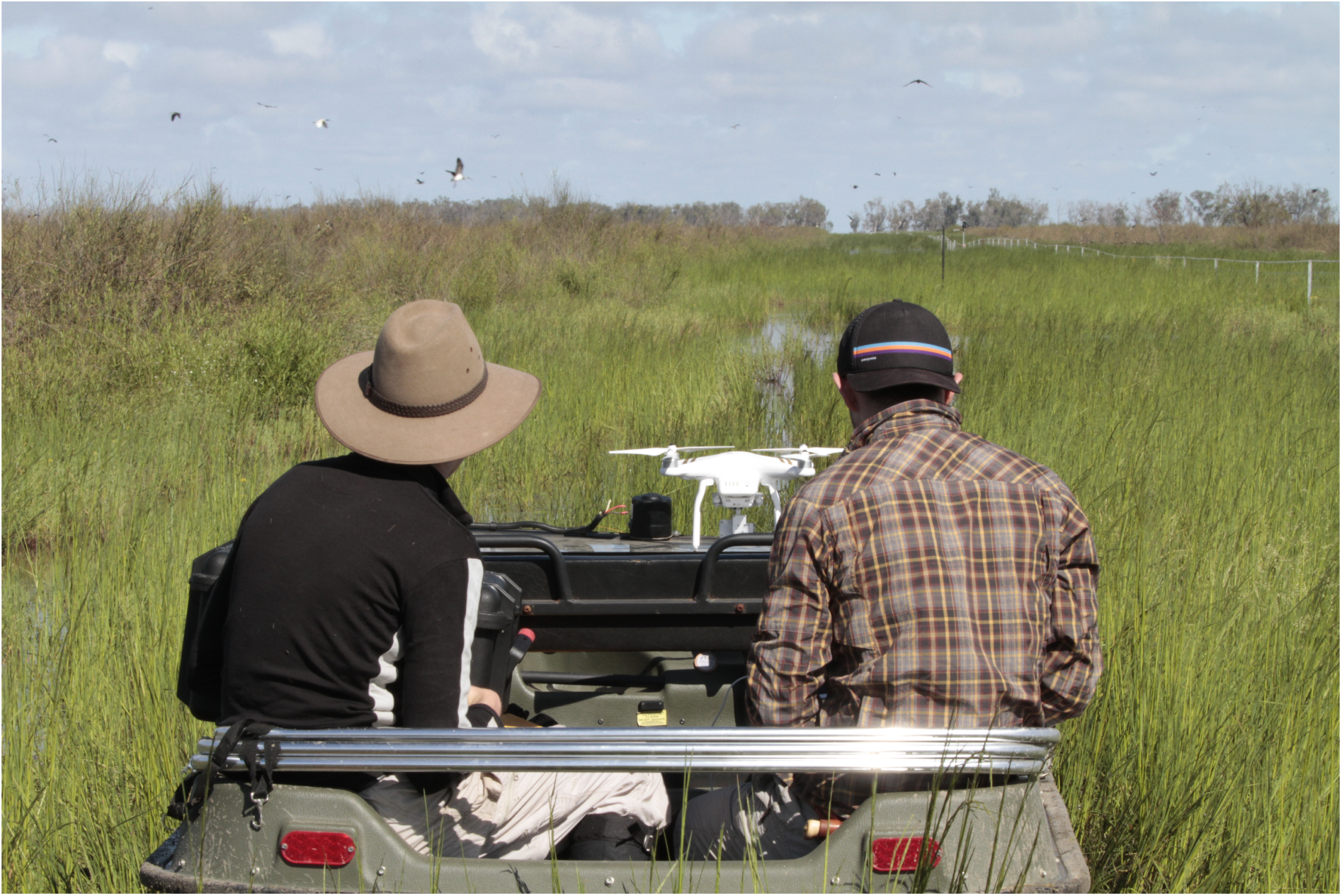
Quad-copter (*DJI Phantom 3 Pro*) launch from an amphibious vehicle (*Argo 8x8 650*) at a straw-necked ibis colony on the Lower Lachlan River in New South Wales, Australia.

### Ibis breeding colonies

The Ibis breeding colonies (Straw-necked Ibis *Threskiornis spinicollis*, Australian White Ibis *T. moluccus*, Glossy Ibis *Plegadis falcinellus*) presented a particularly challenging environment. One of the Lower Lachlan colonies had at least 200,000 adults (100,000 nests) at the time of flying. The other colonies had between 10,000 – 50,000 adults. Ibis usually nest on inundated vegetation inlcuding lignum (*Duma florulenta*) and phragmites (*Phragmites australis*). Nests are typically between 20cm – 2m above ground level. At two of the colonies (Lower Lachlan and Lower Murrumbidgee), we conducted more systematic observations of the impact of the drone on Ibis behaviour, since they were active breeding events. This was in addition to capturing imagery over the entire colony. In order to ensure minimal impact, we monitored the effect of a drone on nesting adults, before conducting the full-colony monitoring exercise. Before any flights had been conducted with the drone, we entered the colony on an amphibious vehicle (*Argo 8x8 650*). After entering the colony, a random group of nests were chosen and a *GoPro Hero 5 Black* fixed to a 2.3 m pole was directed at the nests. We then moved (in the vehicle) approximately 50 m away and out of line of sight of the camera. We waited approximately 20 minutes to allow time for birds to return to their nests before launching the drone. After confirming safe flight parameters, the drone was elevated to 120 m above take off (ATO) and navigated to the nest site being filmed from the ground. The drone was slowly (approx. 1 m/s) descended to 20 m ATO, and hovered for 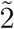 minutes, and then descended to 10 m ATO. The landscape is flat, so height in meters ATO approximates height above the nests. The drone was raised and lowered multiple times at a speed of around 1 m/s to observe the height at which birds flushed from their nests, and under what conditions they returned. The drone was then flown back to the vehicle and we again waited 20 minutes before recovering the GoPro. The drone itself also captured video of the nest sites. Other studies (e.g., Vas et al. (2015)) performed multiple repeated experiments and while this is ideal from an experimental design perspective, we considered any additional disturbance to the birds unnecessary as the subsequent monitoring involved systematic flight lines over the entire colony.

### Animal welfare

The ethics approvals we operated under covered the types of flight patterns for testing interaction with birds, so far as to obtain safe monitoring practices. The ethics requirements explicitly prohibited experimental designs that repeatedly induced interactions (e.g. (Vas et al., 2015)), as it was deemed to cause unnecessary potential risk. This is the primary reason for our relatively *ad hoc* observations.

## Results

### Birds encountered

We encountered a diverse group of bird species across many different environments; some of our sites were over 1500 km apart. In over 70 hours of flights, we had no strikes, nor did we encounter a situation where aggression posed a serious threat. Table details the main birds of interest that we considered might pose a risk at out various field sites. We also provide a list of all other birds observed at each site, that showed no noteworthy interactions with drone operation (Appendix A). Additionally, results from the Ibis colonies are provided in more detail below.

Of most concern in flight planning was the presence of raptors at many of our study sites. However, we did not encounter any negative interactions with raptors. Wedge-tailed Eagles (*Aquila audax*), Australia's largest bird of prey, were common at many of our study sites. At Sturt National Park and Strzelecki Reserve, they were present for the majority of flights, but were not observed to show interest the drone. They were also observed in Yantabulla but were not observed during flight. Black Kites (*Milvus migrans*) and Australian Kestrels (*Falco cenchroides*) were frequently observed at many of our sites outside of the Sydney basin. They appeared quite content to fly in close proximity to the drones, and continued normal activities. For example, while the drone was within 15 meters of an Australian kestrel at one of the Lower Lachlan sites, the kestrel showed no behavioural changes and continued to hunt as normal, resulting in successful prey-capture.

We did observe at least one instance of a negative interaction with the drone, which was from an Australian Magpie (*Cracticus tibicen*) in the Sydney area. During their breeding season, on two occasions (August 2015 and October 2016), they flew aggressively towards the drone, although evasive action by the drone-operator, was effective. In contrast, Pied Currawongs (*Strepera graculina*) left their nests when approached by drones and displayed territorial calls, but not not attempt to physically attack the drone. When Currawongs were similarly approached by other birds (i.e., Channel-billed Cuckoo (*Scythrops novaehollandiae*), Australian Raven (*Corvus coronoides*), Noisy Miner (*Manorina melanocephala*), and Common Myna (*Acridotheres tristis*), they dispalyed both audible and physical territorial behaviour. Moreover, during the non-breeding season, Australian magpies and pied currawongs showed little interest in the drone. Masked Lapwings (*Vanellus miles*) also displayed typical territorial calls, but did not demonstrate any aggressive actions towards drones - masked lapwings nest on open ground, thereby generally minimizing close proximity to the drone. Of minor note is the behaviour of small groups of passerines that were observed within the Sydney basin. Groups of noisy miners and common mynas at times appeared as though they were being aggressive (similar to behavior when a raptor is overhead), but were never observed to strike or attack the drone. Additionally, groups of Welcome Swallows (*Hirundo neoxena*), fairy martins (*Petrochelidon ariel*), and European Starlings (*Sturnus vulgaris*) often flew extremely close (i.e., <1 m) to the drone. On several occasions, swarms of insects were attracted to the multi-rotor drones, though we were uncertain whether these insectivores were attracted to the insects.

### Ibis colonies

Ibis colonies are areas with high densities of nests and birds, meaning adult Ibis were always in close proximity to the drone. This was also true at higher flight altitudes, as Ibis were observed flying in thermals that stretched many 100's of meters into the air. Manual counting of individual nests from the processed drone imagery at one of the Lower Lachlan sites indicated that there were 101,360 nests. Notwithstanding the error associated with that value, which is yet to be fully quantified, it is nonetheless a daunting thought when considering a drone flying operation. We provide (annotated) video of the filmed nest site (*https://youtu.be/86cgvCCcNto*) and we provide a brief summary here. At the Lower Lachlan site, Ibis directly below the drone flushed from their nest when the drone descended to about 20 m. Ibis on adjacent nests (10 to 15 m away) displayed vigilant behavior but did not flush (Fig. 3). If the drone was left hovering at a height of 15 m or greater, birds would return to their nest within 30 seconds to a minute. If the drone was left hovering at 10 m, birds did not return to their nest within 5 minutes, the maximum time we allowed in order to minimise disturbance to chicks and eggs. The flush of birds caused by retrieving the camera (i.e., walking into the colony) was at least 3 to 4 times larger (in number of birds) than that caused by the drone (Fig. 3 and *https://youtu.be/86cgvCCcNto*). Results were almost identical at the Lower Murrumbidgee site, except that birds tended not to flush until the drone descended to between 10-15 m. Ibis occasionally flew quite close to the drone, if they did not see it when changing direction, although they quite easily avoided it. We provide a video of such an avoidance (*https://youtu.be/RQGYJig5-1M*).

**Figure 3:**
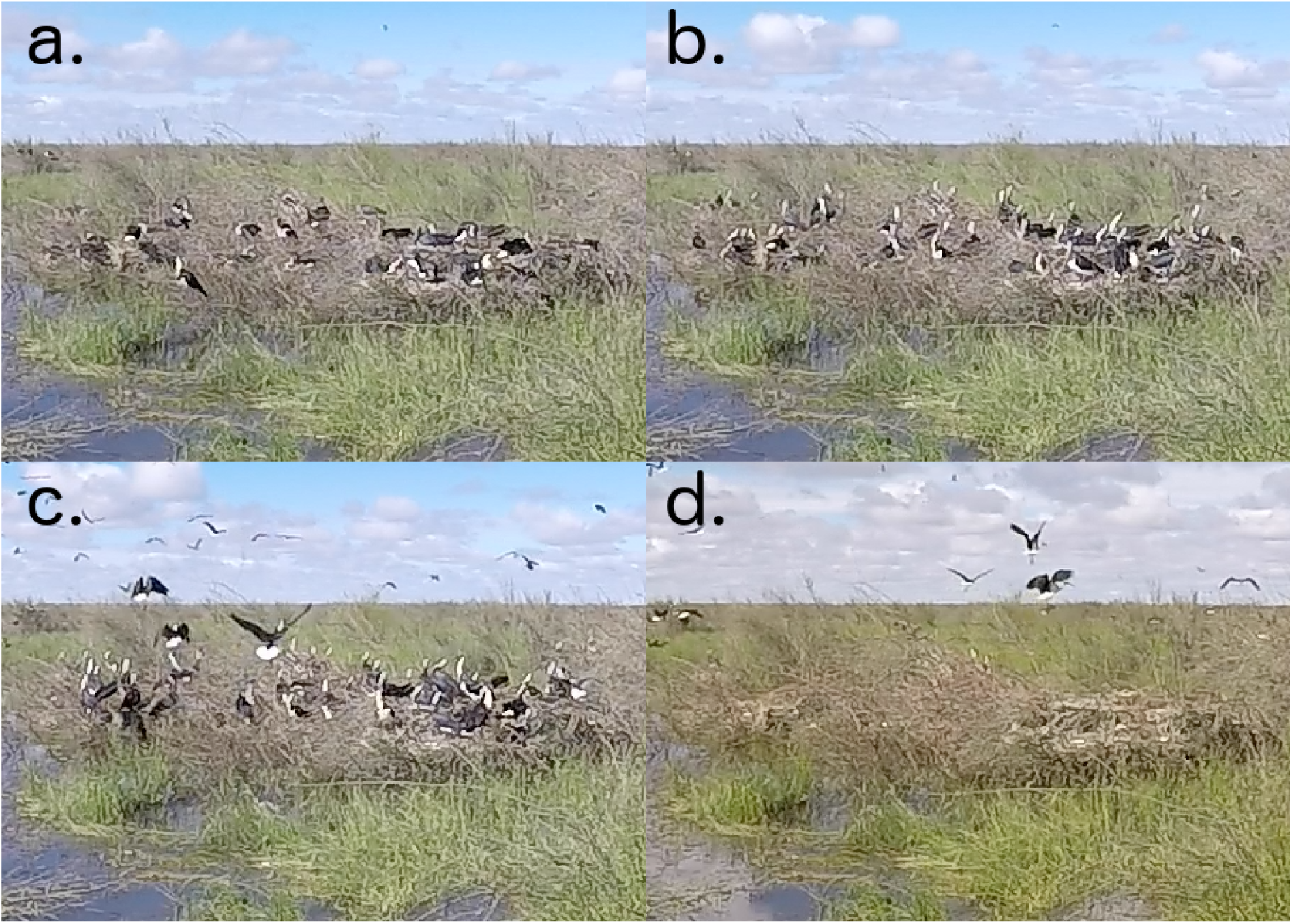
Images of a group of Straw-necked Ibis nests at the Lower Lachlan River in New South Wales, Australia. The nests shown are approximately 15 m away from another group of nests over which a quad-copter drone was being flown. a) shows a typical state pre-disturbance of any kind; b) vigilant behaviour when the drone was lowered to approximately 20 m above the adjacent nests, some birds from the adjacent nests flush; c) more highly vigilant behaviour when the drone was lowered to approximately 10 m above the adjacent nests; d) birds flushed from nest as the camera was retrieved on foot.

## Discussion

Overall, we tended to observe reactions consistent with those reported (or implied) from various drone monitoring studies focused on waterbirds and passerines (Chabot et al., 2015; Descamps et al., 2011; Hodgson et al., 2016; Sardà-Palomera et al., 2012; Vas et al., 2015). Considering this, we think it reasonable that most of the non-territorial birds in Australia are relatively low risk to fly over. We encountered several birds that can be highly territorial and aggressive during breeding season, but only the Australian Magpie showed truly aggressive action towards the drone. Magpies, and to a lesser extent Currawongs and Lapwings, are colloquially bold and will readily harass and strike other birds and people. When Magpies presented a threat we found that an evasive action of flying full speed away, angled upwards, was sufficient to avoid contact. Magpies retreated as per their normal behaviour once the drone was 50-100 m away. Operators should thus always be aware of the breeding season for birds in their study area. There is no existing literature on interactions between drones and raptors, so our findings here provide a basis for further study. Anecdotal evidence suggests that Wedge-tailed Eagles are serious threats to drones, although we did not experience any negative interactions. In fact, none of the raptors present at our sites appeared to be interested in the drones. Large raptors (Wedge-tailed Eagles particularly) tend to be more active in higher winds or during parts of the day where thermals have developed. We avoided those conditions in general, so that may have contributed to the lack of interest shown, and we would certainly encourage others to follow similar guidelines. If a large raptor is observed, we would still recommend safely landing the drone. If a raptor surprises an operator, there is little that can be done except evasive action to land the drone as quickly and safely as possible.

Whilst our work was not systematically designed to test interactions, we show that relatively affordable drones have the capacity for monitoring very large groups of birds, and we feel that maintaining safe flight parameters with relatively low disturbance levels is quite achievable. As far as we know, the Ibis colony at the Lower Lachlan River is the largest bird colony to date to have counts derived from drone imagery. Chabot et al. (2015); Hodgson et al. (2016); Trathan (2004); Descamps et al. (2011) detail monitoring of groups of birds in the order of several thousand to around 11,000. Our work in the Ibis colonies is detailed here to the extent that we think will be useful for others to plan and attempt similar use of drones over large colonies. Further analysis, in context of bird behavior, counting strategies and accuracy, and colony monitoring success, is warranted (Brandis et al., prep). That work will also compare disturbance between drones and traditional monitoring methods, that is, on-foot, canoes, amphibious vehicles and aerial survey. Another major focus for future research is automated processing of the drone imagery products. At present, nest and bird numbers have been manually counted from the imagery, but current research (Lyons et al., prep) is underway that is focusing on automated machine learning and statistical methods. Most current literature is focused on counting bird numbers (Chabot and Francis, 2016), as opposed to counting bird nests, which is often the primary focus for monitoring, particularly in waterbird breeding colonies.

One important aspect we did not measure during our work was the impact of sound. In relatively quiet areas, drones are reasonably noisy, and can be heard 200-300 m away. We are unsure of the impact this is likely to have, and it is likely that the existing literature on the impact of noise on wildlife will turn its attention to drones. Incidentally, while working in the bird colonies, the background noise of the colony was such that the drone was inaudible, to humans, once it was more than 30-40 m away.

In conclusion, we provide considerations to those planning drone monitoring exercises where bird interactions are likely, or where guidance on potential interactions is sought. Firstly, consider carefully the birds likely to be present, if they are territorial, and if they are in breeding season. Start flights by first ascending to a reasonable altitude, as most interactions will occur close to the ground. Raptors are still a risk at higher altitudes, but avoiding the environmental conditions discussed above and having an evasive landing procedure will mitigate that risk. After assessing the area flying at high altitude, lower the drone slowly to obtain an idea of when interactions begin to occur. Needless to say, spotters are invaluable. Although it may seem obvious once stated, there is no need to try and avoid flying birds - they are highly skilled (generally) at avoiding birds in flight. See the video link to ibis avoiding the drone in flight. Additionally, multi-rotor drones are able to come to a complete stop mid-air very quickly; birds typically do not do this, so we recommend avoiding this procedure when operating in close proximity to flying birds. We found that birds tended to become more vigilant or alarmed when the drone was in stationary flight. If a collision is anticipated, then a reduction of pace and change of course are suitable options. This paper adds to the growing literature that highlights the potential of drones for avian research, as well as providing a basis for critical future research to ensure safe and effective monitoring.

**Table 1:**
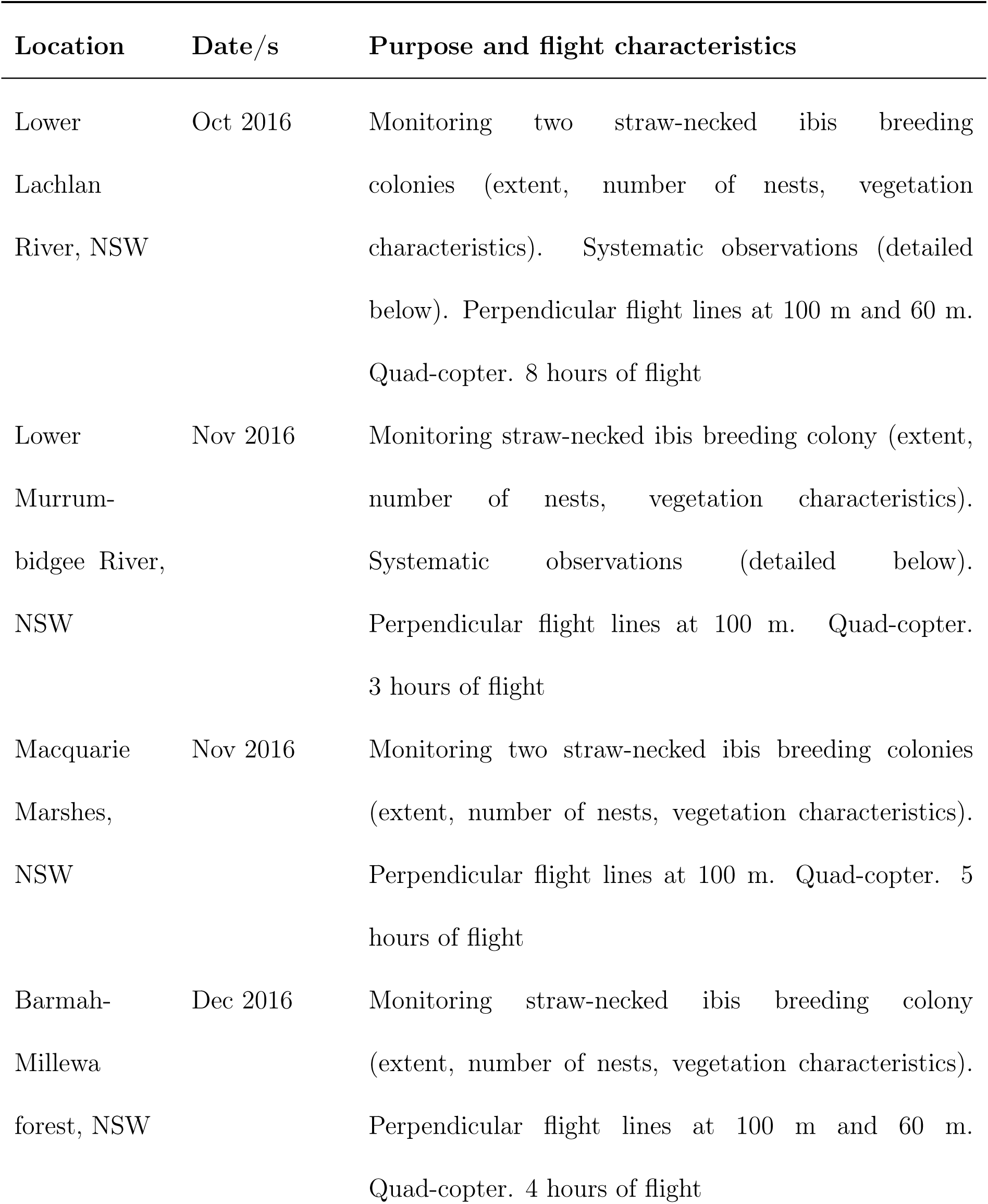
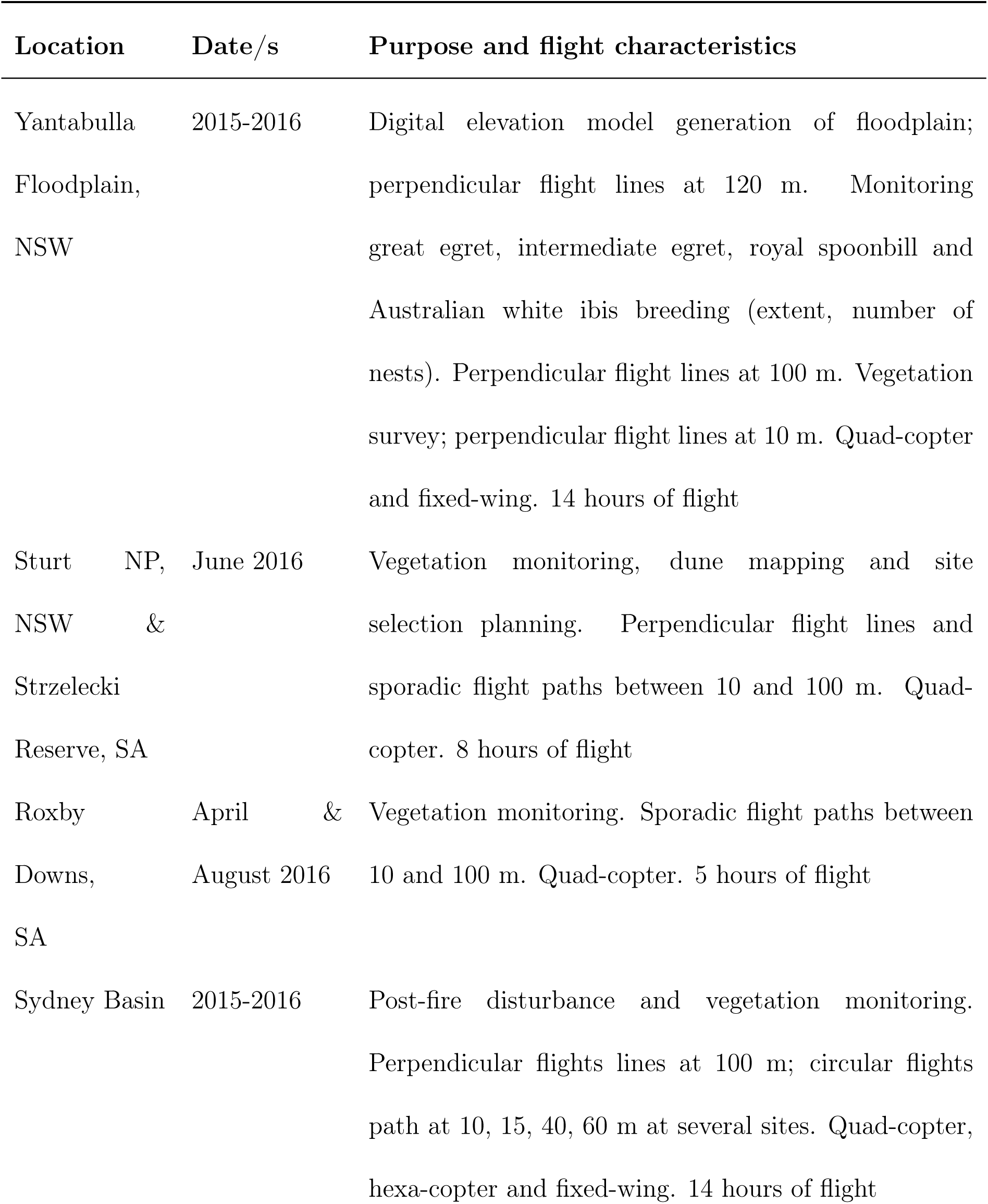
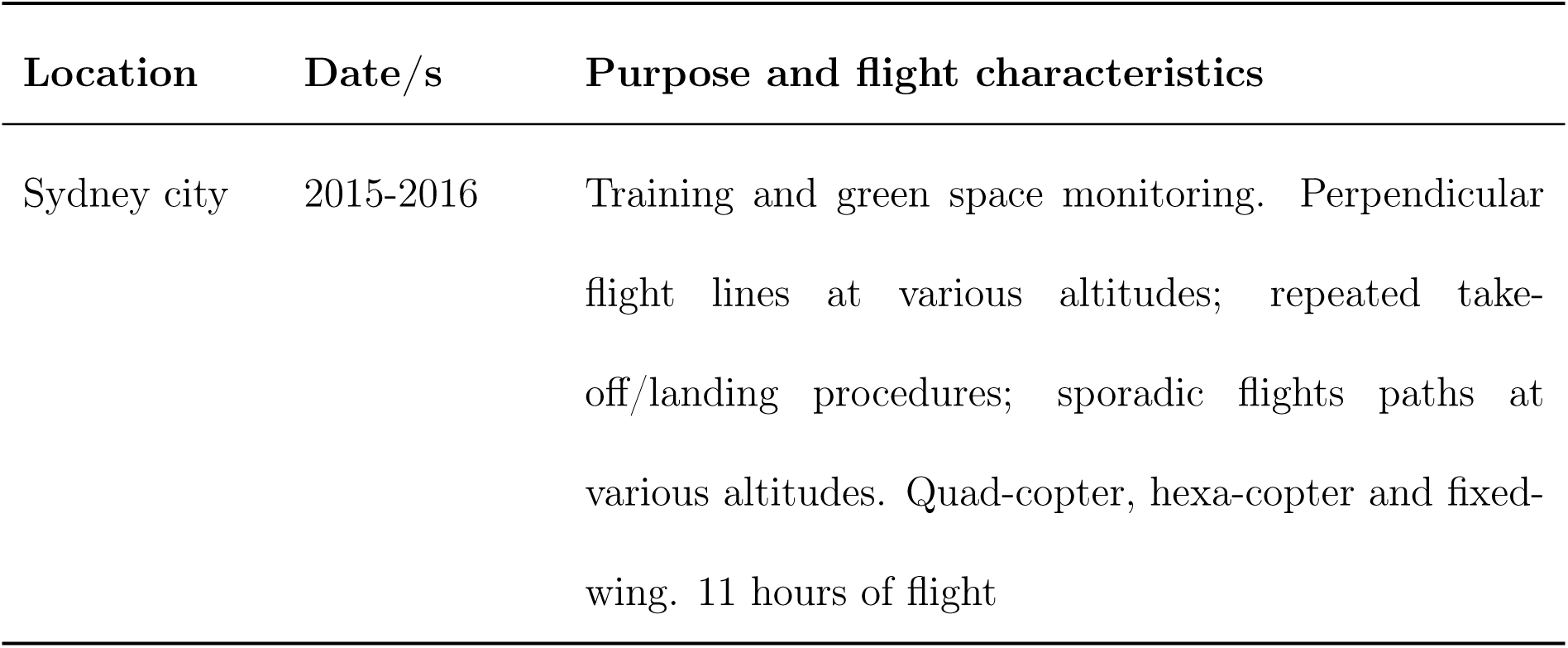
Study site information. All flight heights are above take off (ATO).

| Location | Date/s | Purpose and flight characteristics |
| --- | --- | --- |
| Lower Lachlan River, NSW | Oct 2016 | Monitoring two straw-necked ibis breeding colonies (extent, number of nests, vegetation characteristics). Systematic observations (detailed below). Perpendicular flight lines at 100 m and 60 m. Quad-copter. 8 hours of flight |
| Lower Murrumbidgee River, NSW | Nov 2016 | Monitoring straw-necked ibis breeding colony (extent, number of nests, vegetation characteristics). Systematic observations (detailed below). Perpendicular flight lines at 100 m. Quad-copter. 3 hours of flight |
| Macquarie Marshes, NSW | Nov 2016 | Monitoring two straw-necked ibis breeding colonies (extent, number of nests, vegetation characteristics). Perpendicular flight lines at 100 m. Quad-copter. 5 hours of flight |
| Barmah-Millewa forest, NSW | Dec 2016 | Monitoring straw-necked ibis breeding colony (extent, number of nests, vegetation characteristics). Perpendicular flight lines at 100 m and 60 m. Quad-copter. 4 hours of flight |

| Location | Date/s | Purpose and flight characteristics |
| --- | --- | --- |
| Yantabulla<br>Floodplain,<br>NSW | 2015-2016 | Digital elevation model generation of floodplain; perpendicular flight lines at 120 m. Monitoring great egret, intermediate egret, royal spoonbill and Australian white ibis breeding (extent, number of nests). Perpendicular flight lines at 100 m. Vegetation survey; perpendicular flight lines at 10 m. Quad-copter and fixed-wing. 14 hours of flight |
| Sturt NP,<br>NSW &<br>Strzelecki<br>Reserve, SA | June 2016 | Vegetation monitoring, dune mapping and site selection planning. Perpendicular flight lines and sporadic flight paths between 10 and 100 m. Quad-copter. 8 hours of flight |
| Roxby<br>Downs,<br>SA | April &<br>August 2016 | Vegetation monitoring. Sporadic flight paths between 10 and 100 m. Quad-copter. 5 hours of flight |
| Sydney Basin | 2015-2016 | Post-fire disturbance and vegetation monitoring. Perpendicular flights lines at 100 m; circular flights path at 10, 15, 40, 60 m at several sites. Quad-copter, hexa-copter and fixed-wing. 14 hours of flight |

| Location | Date/s | Purpose and flight characteristics |
| --- | --- | --- |
| Sydney city | 2015-2016 | Training and green space monitoring. Perpendicular flight lines at various altitudes; repeated take-off/landing procedures; sporadic flights paths at various altitudes. Quad-copter, hexa-copter and fixed-wing. 11 hours of flight |

**Table 2:** Key bird species interactions with drones.

| Bird(s) | Sites present | Interactions of note |
| --- | --- | --- |
| Ibis | Lower Lachlan, Lower Murrumbidgee, Barmah-Millewa, and Macquarie Marshes | Present in large numbers, but showed little interest or aversion to drones, except when approached within 10 m |
| Australian Magpie | Coastal and central NSW sites | Aggressive towards drone in breeding season |
| Pied Currawong, Masked Lapwing | Coastal and central NSW sites | Abundant and active during breeding season, but not aggressive towards drone |
| Wedge-tailed Eagle, Black Kite, Whistling Kite, Australian Kestrel | All sites outside Sydney Basin | Observed to be present during many flights, but largely uninterested in drones |
| Waterbirds (ducks, piscivores and waders) | Yantabulla. Non-desert sites outside Sydney Basin | Birds showed no obvious aversion, but tended not to take flight while drone present |

Table 2: Key bird species interactions with drones.
| Bird(s) | Sites present | Interactions of note |
| --- | --- | --- |
| Noisy Miners,<br>Indian Mynas | Sydney Basin sites | At times appear to display aggressive<br>behaviour in close proximity to the drone,<br>but never struck or attacked the drone |
| Swallows,<br>Martins,<br>Starlings | Sydney Basin sites | Groups fly extremely close the drone, but<br>no aggression or contact was observed |

## Acknowledgments

Financial and logistical support from research grants (UNSW grant PS40727 to ML, ARC grant LP150100972), the Commonwealth Environmental Water Office, the New South Wales Office of Environment and Heritage, Bush Heritage Australia, Arid Recovery Reserve and local land owners. We operated under two animal ethics approvals from the University of New South Wales Animal Care and Ethics committee (approval numbers 16/3B and 16/131B).

### Appendix A

This appendix provides a list of birds observed at each study location during drone flying operations, that are not directly discussed (or are mentioned in their broader taxonomic group) in the main text and showed no notable interaction with the drones. Some study sites were relatively small, or had more limited survey, meaning less birds were observed. Section and Table in the main text provides further information about the study locations.

#### Lower Lachlan River

Plumed Whistling-Duck, Black Swan, Pacific Black Duck, Grey Teal, Pink-eared Duck, Hardhead, Hoary-headed Grebe, Little Pied Cormorant, Australian Pelican, Great Egret, Glossy Ibis, Australian White Ibis, Royal Spoonbill, Swamp Harrier, Black Kite, Whistling Kite, Australasian Swamphen, Eurasian Coot, Pied Stilt, Whiskered Tern, Crested Pigeon, Galah, Superb Fairywren, Magpie-lark, Australian Reed-Warbler, Little Grassbird

#### Lower Murrumbidgee River

No additional birds observed

#### Macquarie Marshes

Royal Spoonbill

#### Barmah-Millewa Forest

Royal Spoonbill, Great Egret, White-faced Heron, Musk Duck, Australasian Swamphen

#### Yantabulla Floodplain

Great Egret, Intermediate Egret, Australian White Ibis, Yellow-billed Spoonbill, Royal Spoonbill, Australian Pelican, Australian Wood Duck, Pacific Black Duck, Grey Teal, Pink-eared Duck, Little Pied Cormorant, Australasian Darter, White-necked Heron, White- faced Heron, Eurasian Coot, Pied Stilt, Black-fronted Dotterel, Peaceful Dove, Sacred Kingfisher, Cockatiel, White-plumed Honeyeater, Willie Wagtail, Magpie-lark

#### Sturt National Park and Strzelecki Reserve

White-winged Fairy-wren, Masked Woodswallow, Singing Honeyeater, Black-faced Woodswallow, Zebra Finch, Cinnamon Quail-Thrush, Chirruping Wedgebill

#### Roxby Downs

Black-faced Woodswallow, Crested Pigeon, Little Raven, Zebra Finch

#### Sydney Basin

Yellow-faced Honeyeater, Eastern Spinebill, Red Wattlebird, Noisy Friarbird, New Holland Honeyeater, Gray Butcherbird, Maned Duck, Pacific Black Duck

#### Sydney City

Rainbow Lorikeet, Black-faced Cuckooshrike, Common Koel, Little Corella, Sulphur- crested Cockatoo, Galah, Gray Butcherbird

